# Formin3 regulates dendritic architecture via microtubule stabilization and is required for somatosensory nociceptive behavior

**DOI:** 10.1101/227348

**Authors:** Ravi Das, Jamin M. Letcher, Jenna M. Harris, Istvan Foldi, Sumit Nanda, Hansley M. Bobo, József Mihály, Giorgio A. Ascoli, Daniel N. Cox

## Abstract

The acquisition, maintenance and modulation of dendritic architecture are critical to neuronal form, plasticity and function. Morphologically, dendritic shape impacts functional connectivity and is largely mediated by organization and dynamics of cytoskeletal fibers that provide the underlying scaffold and tracks for intracellular trafficking. Identifying molecular factors that regulate dendritic cytoskeletal architecture is therefore important in understanding mechanistic links between cytoskeletal organization and neuronal function. In a neurogenomic-driven genetic screen of cytoskeletal regulatory molecules, we identified Formin3 (Form3) as a critical regulator of cytoskeletal architecture in *Drosophila* nociceptive sensory neurons. Form3 is a member of the conserved Formin family of multi-functional cytoskeletal regulators and time course analyses reveal Form3 is cell-autonomously required for maintenance of complex dendritic arbors. Cytoskeletal imaging demonstrates *form3* mutants exhibit a specific destabilization of the dendritic microtubule (MT) cytoskeleton, together with defective dendritic trafficking of mitochondria, satellite Golgi and the TRPA channel Painless. Biochemical studies reveal Form3 directly interacts with MTs via FH1-FH2 domains and promotes MT stabilization via acetylation. Neurologically, mutations in human Inverted Formin 2 (*INF2;* ortholog of *form3*) have been causally linked to Charcot-Marie-Tooth (CMT) disease. CMT sensory neuropathies lead to impaired peripheral sensitivity. Defects in *form3* function in nociceptive neurons results in a severe impairment in noxious heat evoked behaviors. Expression of the INF2 FH1-FH2 domains rescues *form3* defects in MT stabilization and nocifensive behavior revealing conserved functions in regulating the cytoskeleton and sensory behavior thereby providing novel mechanistic insights into potential etiologies of CMT sensory neuropathies.

**Significance Statement:** Mechanisms governing cytoskeletal architecture are critical in regulating neural function as aberrations are linked to a broad spectrum of neurological and neurocognitive disorders. Formins are important cytoskeletal regulators however their mechanistic roles in neuronal architecture are poorly understood. We demonstrate mutations in *Drosophila formin3* lead to progressive destabilization of the dendritic microtubule cytoskeleton resulting in severely reduced arborization coupled to impaired organelle and ion channel trafficking, as well as nociceptive sensitivity. INF2 mutations are implicated in CMT sensory neuropathies, and INF2 expression can rescue microtubule and nociceptive behavioral defects in *form3* mutants. While CMT sensory neuropathies have been linked to defects in axonal development and myelination, our studies connect dendritic cytoskeletal defects with peripheral insensitivity suggesting possible alternative etiological bases.

## Introduction

Dendrites function in the reception, integration and propagation of sensory or synaptic information and thereby mediate neural computation. Specification and maturation of diverse dendritic architectures is subject to complex intrinsic and extrinsic regulatory programs that modulate the formation and plasticity of a functional nervous system (Lefebvre et al. 2015). An important convergent regulatory target of extrinsic signaling and intrinsic factors is modulation of the neuronal cytoskeleton, which ultimately contributes to dendritic form and function (Das et al. 2017; Nanda et al. 2017). Thus, elucidating molecular bases underlying dendritic cytoskeletal organization is crucial to uncovering principles governing neuronal diversity, as well as potential mechanistic links between architecture and behavior.

Regulation of the cytoskeleton is critically important to the establishment, maintenance, and plasticity of neural morphology, as well as function (Franker and Hoogenraad 2013). Upon achieving a mature shape, dendritic arbors retain a certain degree of plasticity that occurs via dynamic cytoskeletal modulation, thereby contributing to refinements in neural connectivity. Moreover, stabilization of mature dendrites is required for proper neuronal function, as morphological destabilization can lead to functional impairments in neurotransmission, result in neurodegeneration, disrupt organelle trafficking, and in the case of sensory neurons, disrupt responses to environmental stimuli leading to behavioral defects such as peripheral insensitivity (Nanda et al. 2017; Honjo et al. 2016). Architectural organization and dynamic modulation of cytoskeletal fibers is controlled by a vast array of regulatory factors that impact assembly, disassembly, bundling, severing, stabilization and motor-based transport (Coles and Bradke 2015; Kapitein and Hoogenraad 2015). In addition, defects in cytoskeletal-mediated motor-based transport have been shown to result in loss of dendritic identity, defects in neuronal polarity, and severe impairments in dendritic development (Rolls 2011; Nanda et al. 2017).

Formins represent one important family of evolutionarily conserved, multi-functional cytoskeletal regulators (Goode and Eck 2007). Formins are multidomain, dimeric functional molecules characterized by conserved Formin Homology (FH) domains (Higgs 2005). The functional role of Formins in actin polymerization has been well documented whereby the FH2 domain initiates actin nucleation and remains bound at the barbed end of an F-actin filament moving along the growing filament promoting elongation by preventing access of capping proteins (Chhabra and Higgs 2007). F-actin elongation is further enhanced by association of Profilin with the Formin FH1 domains (Kovar 2006). Unlike other actin nucleators, such as Arp2/3, that form branched actin filaments, Formins assemble linear actin filaments (Evangelista et al. 2002). Despite their basic function and properties, Formins vary significantly, for instance, some bundle actin filaments, some sever or depolymerize actin filaments, and in recent studies, Formins have been implicated in regulating microtubule (MT) organization and dynamics (Bartolini and Gundersen 2010; Breitsprecher and Goode 2013; Roth-Johnson et al. 2014). Formins appear to stabilize MTs both through their direct binding and/or by altering their post-translational state (Gaillard et al. 2011; Thurston et al. 2012). However, little is known regarding the potential functional roles of Formins in neural development and dendrite arborization.

Here, we provide mechanistic links between Formin-mediated dendritic cytoskeletal stabilization and nociceptive behavioral sensitivity. Specifically, we demonstrate that Formin3 (Form3) is specifically required for dendritic arbor maintenance, which occurs predominantly via roles in MT stabilization. MT destabilization is associated with defective dendritic trafficking of mitochondria and satellite Golgi, as well as Golgi fragmentation. Form3 directly interacts with MTs via FH1-FH2 domains and promotes MT acetylation. Intriguingly, defects in *INF2* have been causally linked to CMT sensory neuropathies known to cause impaired peripheral sensitivity. Disruption of *form3* in nociceptive neurons severely impairs noxious heat evoked behaviors and leads to defects in dendritic trafficking of the TRPA channel Painless. Moreover, *form3* mutant defects in nociceptive behavioral sensitivity and dendritic MT stabilization can be rescued by expression of INF2 FH1-FH2 domains. Collectively, our findings suggest *form3* and *INF2* share conserved functions in regulating the dendritic cytoskeletal and sensory behavior providing new mechanistic insights into potential etiologies underlying CMT sensory neuropathies.

## Materials and methods

### Drosophila husbandry and stocks

*Drosophila* stocks were maintained at 25°C and genetic crosses were performed at 29°C. The following stocks used in this study where ‘B’ represents stocks from Bloomington Stock Center and ‘v’ represents Vienna Stock center, and additional sources are provided parenthetically: *GAL4^477^,UAS-mCD8::GFP/Cyü,tuhP-GÄL·80;GÄL·4^ppk.1.9^,UAS-mCD8::GFP* (aka *CIV-GAL4*), *GAL4^477^;ppk::tdTomato*, *UAS-GMA::GFP;GAL4^477^,UAS-Jupiter::mCherry* (Das et al. 2017), *UAS-form3-B1* (Tanaka et al. 2004), *form3^Em41^,FRT^2A^* (Tanaka et al. 2004, this study), *form3^Em31^,FRT^2A^* (Tanaka et al. 2004, this study), *Df(3L)Exel6110, GAL4^¦40^,UAS-Venus,SüP-FLP^42^; +; tuhP-GAL80,FRf^A^* (DGRC stock 109-950), *UAS-mito-HA-GFP.AP* (B8442), *UAS-manII::EGFP, GAL4^ppk^,UAS-myr::GFP, UAS-INF2-FH1-FH2*, *UAS-INF2-Full* length, *GAL4^ppk^, UAS-ChETA-YFP, UAS-Painless^p103^::VFP* (Hwang et al. 2012), *UAS-capu-IR* (B2922), *UAS-dia-IR* (B33424, B35479), *UAS-Fhos-IR* (B31400, B51391), *UAS-Frl-IR* (B32447, v110438), *UAS-DAAM-IR* (B39058), *UAS-form3-IR* (B32398 (focus of analyses in this study), v45594, v42302, v28437, v107473), *UAS-exo70-IR* (B55234, B28041), *UAS-sec8-IR* (B57441), *UAS-CG3967-IR* (B28777, v26338, v26339, v106247), *UAS-CG17003-IR* (v32804, v32805, v101273) and *UAS-mtrm-IR* (B35430). *OregonR (ORR)* served as a wild-type strain.

### IHC analysis and live confocal imaging

Larval filets were processed and labeled as described (Sulkowski et al. 2011). Primary antibodies used in this study include: rabbit anti-Form3 (1:100), mouse anti-Futsch (22C10) (1:200; DSHB), anti-HRP (1:200; Jackson Immunoresearch), mouse anti-acetylated α Tubulin (6-11B-1) (1:100; Santa Cruz sc-23950), chicken anti-GFP (1:1000; Abcam). Donkey anti-rabbit, anti-mouse and anti-chicken secondary antibodies (Jackson Immunoresearch) were used at 1:200. Anti-Form3 polyclonal antibody was generated in rabbits using KLH-conjugated peptides (GenScript). Form3 epitope (RGSDASSPTRKPSQ (amino acids (aa) 321-334)) was predicted by the GenScript OptimumAntigen design tool. Anti-Form3 antibody was affinity purified against antigenic peptide and specificity confirmed via IHC analyses. IHC slides were then mounted in Fluoromount Aqueous Mounting Medium (Sigma), and imaged at room temperature on a Zeiss LSM780 confocal system with either a 20× (dry), 40 × or 60 × (oil immersion) objective. Zen blue software was used to quantify the mean intensity of the fluorescence. Live confocal imaging was performed as described (Iyer et al. 2013). Briefly, 6–10 third instar larvae were analyzed for neurometric quantitation from segments A3-A5. For time course analyses, larvae were live imaged at the indicated time point, returned to agar plates, aged at 25°C, and then re-imaged. Images were collected as z-stacks with a step-size of 1.0-2.0μm and 1024×1024 resolution.

### MARCM analysis

MARCM analyses were performed as described (Sulkowski et al. 2011). Briefly, *form3^Em31^*,*FRT^2A^* or*form3^Em41^*,*FRT^2A^* flies were crossed to *GAL4^5-40^,UAS-Venus,SOP-FLP^42^;* +*; tubP-GAL80,FRT^2A^* flies (DGRC stock 109-950). Third instar larvae with GFP-labeled *form3* mutant neurons were subjected to live confocal microscopy.

### Neuromorphometric quantification and multichannel reconstructions

Quantitative neurometric analyses were performed as previously described (Das et al. 2017). Maximum intensity projections of z-stacks were exported using Zen-blue software. Exported images were manually curated to eliminate non-specific auto-fluorescent spots such as larval denticle belts using a custom designed program, *Flyboys*. Images were processed, skeletonized and neurometric data compiled as described (Iyer et al. 2013). Quantitative neurometric information was extracted and compiled using custom Python algorithms freely available upon request. The custom Python scripts were used to compile the output data from the Analyze Skeleton ImageJ plugin and the compiled output data was imported into Excel (Microsoft). Neurometric data was analyzed in Microsoft Excel and statistical tests were performed and plotted in GraphPad Prism 7. For Sholl analysis, we used NeuronStudio (Wearne et al. 2005) to plot the density profiles of branches as a function of distance from the cell soma; to determine the peak of maximum branch density (critical value/ # of intersection) and its corresponding radius. For Strahler analysis, we used the centripetal labeling function of NeuronStudio. (Wearne et al. 2005). Dendritic terminals are defined as Strahler order 1. Order increases when 2 or more branches of the same order intersect at a branching point. For Strahler analysis for Vaa3D we used L-Measure (Scorcioni et al. 2008).

Multichannel cytoskeletal reconstructions and quantitative analyses were performed as previously described (Das et al. 2017). Briefly, two channel (GFP for F-actin and RFP for MT) image stacks (.czi file format) were first processed in Fiji (Schindelin et al. 2012) where a third pseudo-channel was created by adding the signals from the two original channels. These new files with three channels were then imported to Vaa3D (Peng et al. 2014), and the third pseudo-channel was manually reconstructed into the SWC file format (Cannon et al. 1998). These initial traced SWC file and the image stacks were then reopened in Neutube (Feng et al. 2015), and additional tracing, editing and quality checks were conducted. Remaining topological errors were programmatically repaired in batch, by building small custom scripts within the TREES toolbox (Cuntz et al. 2010) package in the MATLAB environment (MathWorks, Natick, MA). The corrected reconstruction files and the image stacks were used as input in Vaa3D plugin to create multichannel ESWC files that represent the morphology along with the intensity and volume occupied by each channel. Then the internal and external structural features were quantified using L-Measure (Scorcioni et al. 2008). Reversed Strahler order based cytoskeletal quantification of ESWC files was carried out in a new analysis program built using TREES toolbox functions (Cuntz et al. 2010). MT or F-actin quantity of a compartment is defined as (the relative signal intensity of the compartment) * (the volume of the compartment) * (the fraction of the volume occupied by the MT or F-actin signal). Total relative quantity of MT or F-actin has been quantified against path distance from the soma at 40 micron intervals (binning) and is thus not the average relative quantities per unit length (i.e. not normalized to length). As such, smaller vs. larger total arbor length does not influence these quantitative comparative analyses (Das et al. 2017).

### Biochemical assays

Coding sequences of the Form3 FH1-FH2 and FH2 domains were cloned in *EcoRI* cut pGEX2T vector. The resulting plasmids were used to express FH1-FH2 and FH2 only fragments of Form3 as GST fusion proteins in BL21 *E. coli*. Protein purification was carried out as previously described (Barkó et al. 2010) with some minor modifications. For MT binding assays, MTs were assembled from tubulin protein (Cytoskeleton, Inc.) according to the vendor’s instructions. In MT co-sedimentation assays, GST::Form3-FH1-FH2 and GST::Form3-FH2 proteins were pre-cleared by ultracentrifugation, then they were diluted in MT binding-buffer (MBB; 10 mM Na-HEPES pH: 7.0, 1 mM MgCl2, 1 mM EGTA, 1 mM DTT, 20 uM taxol, 0.5 mM thesit, 10 % glycerol) and mixed with pre-assembled MTs (0.5 μM). Control samples did not contain MTs. Protein mixtures were incubated for 30 min at room temperature then centrifuged at 100,000 X g for 1 h at 25 °C. Proteins in the supernatants and pellets were resolved by SDS-PAGE then stained with colloidal Coomassie-blue. In GST pull-down assays, purified Form3 proteins were immobilized on glutathione-S-sepharose beads, followed by incubation with taxol-stabilized MTs (0.5 μM) for 30 min at room temperature in MBB. The beads were washed thoroughly in MBB followed by protein elution in SDS-PAGE sample buffer and detection by anti-GST (Sigma) and anti-α-tubulin (Sigma) Western blot.

### Generation of human INF2 rescue transgenes

For optimal expression, we synthesized *D. melanogaster-codon-optimized* INF2 cDNAs (GenScript). Two custom gene syntheses were performed to generate a full-length cDNA and a cDNA in which the DID and DAD autoinhibitory regulatory domains have been deleted leaving only the FH1 and FH2 domains (FH1-FH2). Each synthesized gene was C-terminally FLAG-tagged (DYKDDDDK) and subcloned into*pUAST-attB*. Transgenic INF2 strains were generated by ΦC31-mediated integration with targeting to 2L (*attP40*) (GenetiVision).

### Behavioral and optogenetic assays

For noxious heat nociceptive assays, age-matched third instar larvae were recovered and briefly rinsed with water to remove any residual media. Larvae were then transferred to black aluminum metal plate which was pre-sprayed with water to generate a thin film facilitating larval movement. Larvae were allowed to acclimate to the plate and resume normal peristaltic locomotion before the plate was transferred to a temperature controlled Peltier plate (TE Technology). The temperature was preset to 45°C to evoke nocifensive behaviors. Heat evoked behaviors were recorded using a Nikon D5300 DSLR camera. Video files were processed using ImageJ and manually curated to evaluate response latency and characterize nocifensive behaviotypes (Chattopadhyay et al. 2012). The maximal latency period for behavioral response was set at 20 sec after stimulus and larvae that failed to exhibit a behavioral response in this interval were classified as non-responders. Optogenetic assays were performed essentially as previously described (Turner et al. 2016). Briefly, third instar larvae expressing CIV>*UAS-ChETA-YFP* were subjected to optogenetic activation (480 nm) and behavioral response recorded. Larvae were reared on media ±1 mM all *trans*-retinal. For optogenetic stimulation, larvae were suspended in a hanging water droplet and exposed to blue light in 10-30 sec intervals. Larvae executing at least one full 360° roll in that interval were scored as responders.

### Statistics and data availability

Data is reported as mean and error bars represent standard error of the mean (SEM) or standard error of proportion (SEP) as indicated in figure legends. Data represent biological replicates with the exception of the Western blots for the GST pull-down assays which represent technical replicates. Statistical analyses were performed using either one-way ANOVA with Bonferroni correction for multiple comparisons; two-way ANOVA with Bonferroni correction; or unpaired t-test. All datasets were tested for normality (Shapiro-Wilk normality test) and homogeneity of variance (Bartlett’s test or F test) before statistical analysis. All new genotypes presented here are available upon request. Digital reconstructions of neuronal morphology have been deposited into the NeuroMorpho.Org database (Ascoli 2006) for public distribution under the Cox and Ascoli archives.

## Results

### form3 regulates dendritic arborization in Drosophila nociceptive sensory neurons

Molecular control of cytoskeletal organization and dynamics is essential in specifying, stabilizing and modulating dendritic architecture. To gain insight into this process, we previously conducted a systematic neurogenomic-driven genetic screen to identify putative cytoskeleton regulators of dendritic arborization using *Drosophila* Class IV (CIV) nociceptive sensory neurons as a model system (Das et al. 2017). Among many genes uncovered, we identified multiple members of the Formin gene family, including *form3* and *Frl* (Das et al. 2017). Formins have been demonstrated to function by regulating the F-actin and microtubule cytoskeletons, however, their potential role(s) in neuronal development, and more specifically dendritogenesis, are poorly understood. Therefore, we performed a pilot RNAi screen to examine potential functional roles of the six *Drosophila* Formins. Intriguingly, our screen revealed that only *form3* knockdown (*form3-IR*) produced defects in CIV dendritic arborization (Figure 1-1A-H). Based upon these results, we chose to conduct more in depth phenotypic studies of *form3* function in dendritic morphogenesis.

**Figure 1:**
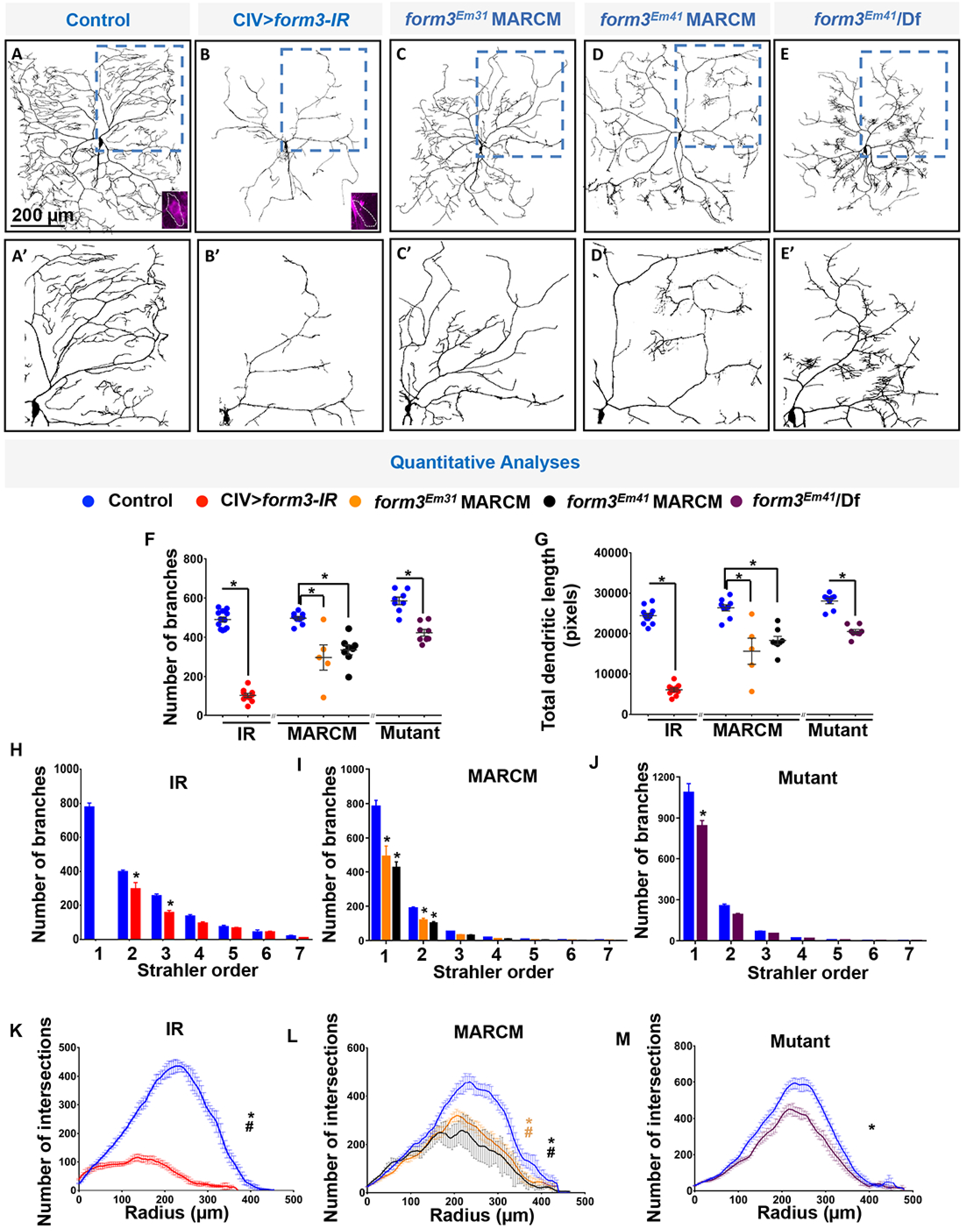
form3 is required for dendritic growth and higher order branching complexity. Representative images of dendritic arborization in CIV (ddaC) neurons in (**A**) control (N=12 neurons) (**B**) *form3-IR* (IR) (N=10 neurons); (**C**) *form3^Em31^* MARCM (N=5 neurons); (**D**) *form3^Em41^* MARCM (N=8 neurons); (**E**) *form3^Em31^/Df* (N=8 neurons). (**A,B**) Insets show CIV soma labeled for Form3 protein expression revealing specificity of antibody and efficacy of *form3-IR* knockdown. (**F,G**) Quantitative dendritic analyses. Values are the mean ± SEM. (**H-J**) Reverse Strahler analysis of control vs. *form3-IR* (**H**), MARCM (**I**) and mutant/Df (**J**). Values are the mean ±SEM for the number of dendritic branches in each branch order (Strahler order) where 7=primary branch from cell body and 1=terminal branches. (**G**) Sholl profile of control vs IR (**K**), MARCM (**L**) and mutant/Df (**M**). Values are the mean ±SEM for the number of intersections as a function of radius distance from the cell body (zero) (Sholl). Statistical tests performed in: (**F_IR_**) unpaired t-test p=0.0001, (**F_MARCM_**) one-way ANOVA with Bonferroni correction p=0.0005 (Em31) and p=0.0011 (Em41), (F_Mutant_) unpaired t-test p=0.00002; (Gir) unpaired t-test p=0.00001, (**G_MARCM_**) one-way ANOVA with Bonferroni correction p=0.0003 (Em31) and p=0.0011 (Em41), (**G_Mutant_**) unpaired t-test p=0.000001. (**H, I, J**) two-way ANOVA with Bonferroni correction: (**H_2,3_**) p=0.0001; (**I_1-Em31,Em41_**) p=0.0001, (**I_2-Em31_**) p=0.0119, (**I_2-Em41_**) p=0.0005; (**J_1_**) p=0.0001. (**K**) unpaired t-test for critical value (*) and the corresponding radius (#) p=0.0001 for the both parameters. (**L**) one-way ANOVA with Bonferroni correction for critical value (*) and the corresponding radius (#), p=0.01469 (Em31 *), p=0.04517 (Em31 - #), p=0.00001 (Em41 *), p=0.02192 (Em41 #); (**M**) unpaired t-test for critical value (*) where p=0.00286

**Extended Figure 1-1:**
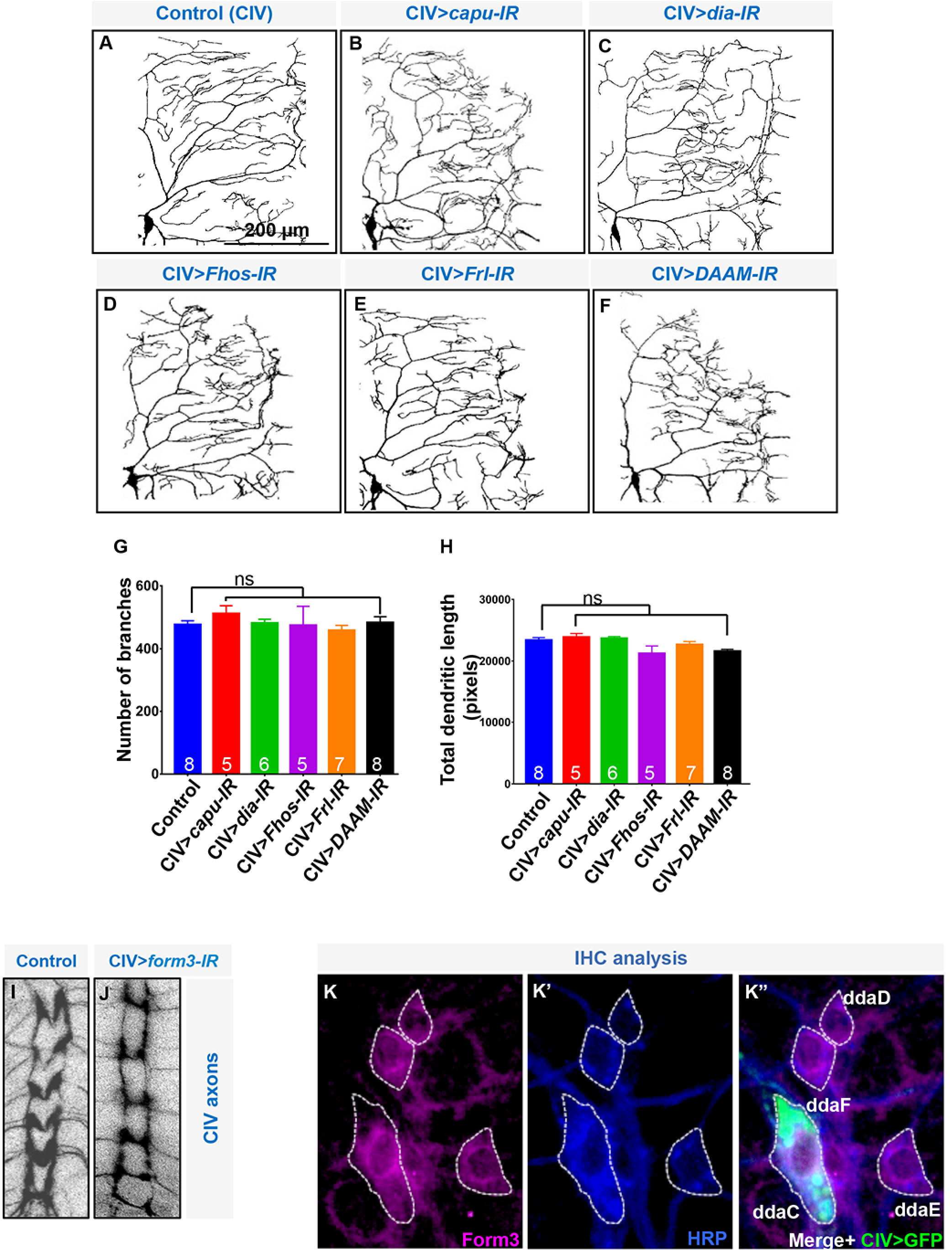
Phenotypic analyses of Drosophila Formins. (**A-F**) Representative images of CIV dendrites in control (**A**) and (**B-F**) RNAi knockdowns for *Drosophila* Formins: *capu; dia; Fhos; Frl; DAAM* (**B-F**). (**G,H**) Quantitative neuromorphometric analyses (mean ± SEM) with N value indicated on the graphs. (**I,J**) Representative images of CIV axonal patterning in the VNC (N=8 of each genotype). (**K-K”**) Class IV-specific *GAL4^477^,UAS-mCD8::GFP* third instar larval filets triple labeled with Form3 (**K**), HRP (**K’**), and Merge+GFP (**K”**). Representative image of N=8 dorsal da neuron clusters. Statistical tests performed (**G,H**): One way ANOVA with Bonferroni correction; p>0.05.

CIV-specific *form3-IR* expression led to severe dendritic reductions resulting in a highly rudimentary arbor with pronounced defects in distal higher order branching relative to controls (Figure 1A,B,F,G,H,K). Dendritic branch order analyses showed significant reductions in higher branch orders (Strahler order 2, 3) compared to control, while Strahler order 1 branches, which represent terminals accounting for the majority of CIV branches, were undetectable in *form3-IR* neurons (Figure 1H). To quantify effects on dendritic branch distribution, Sholl analyses were used to plot the density of profiles of branches as a function of distance from the soma and compare the peak of maximum branch density (critical value) and its corresponding radius. Both parameters were dramatically reduced in CIV-specific *form3-IR* expression relative to controls (Figure 1K).

To independently validate phenotypic defects observed with *form3-IR*, we conducted MARCM clonal analyses using two previously published *form3* point mutant alleles, namely *form3^Em31^* and *form3^Em41^* (Tanaka et al. 2004). Consistent with *form3-IR* analyses, MARCM mutant clones for these alleles revealed defects in CIV dendritic arborization characterized by reductions in distal terminal branching and short interstitial branches (Figure 1C,D). Morphometric analyses revealed significant reductions in both total dendritic branches and concomitant reductions in total dendritic length compared to controls (Figure 1F,G). Moreover, Strahler analysis showed a similar result to that was observed with *form3-IR*, where the reduction was observed predominantly on the terminal branches (Figure 1I). Sholl analyses identified a significant reduction in the critical value for both *form3^Em31^* and *form3^Em41^* (Figure 1L). In addition, we also examined the *form3^Em31^* mutant allele over a chromosomal deficiency (Df) for *form3*. This genetic background produced a phenotype similar to *form3^Em31^* MARCM clones, however there was a notable increase in short, clustered interstitial branches (Figure 1E). This phenotypic difference may be attributable to the heterozygous deletion of one or more of the other eight genes completely deleted by this chromosomal deficiency. Quantitatively, *form3^Em31^/Df* also led to a decrease in both number of dendritic branches and total dendritic length (Figure 1F,G). Likewise, we observed reductions in terminal branches shown via Strahler analysis (Figure 1J) as well as a modest, but significant, reduction in the critical value revealed by Sholl analysis (Figure 1M).

To determine whether *form3* exerts effects specifically on dendrites vs. axons, we also examined overall CIV axon projections and ventral nerve cord (VNC) patterning of axon terminals. Phenotypic analyses revealed a modest qualitative reduction in thickness of select commissural axon projections; however, the longitudinal fascicles appeared largely normal (Figure 1-1I,J). However, given that no gross morphological defects in CIV axon path finding or patterning of axon terminals in the VNC were observed, these results suggest Form3 has a compartment-specific role in regulating dendritic development, though we cannot explicitly rule out subtle defects at individual CIV axon terminals.

To investigate Form3 protein expression, we developed polyclonal antibodies and performed immunohistochemistry (IHC) on fileted third instar larvae revealing Form3 is expressed in da neuron subclasses including CIV neurons (Figure 1-1K-K”). To assess antibody specificity and validate *form3-IR* mediated knockdown, we examined Form3 labeling upon CIV-specific expression. Relative to controls, these analyses revealed a strong reduction in Form3 protein levels in CIV neurons (Figure 1A,B bottom right insets).

Collectively, these data indicate that Form3 promote the appropriate number and positions of branches along the proximal-distal axis of dendritic arbors, and are required to promote higher order branches. Given that *form3-IR* produced the strongest phenotypic defects on CIV dendritogenesis, and that the *form3* homozygous mutant alleles are lethal prior to the third instar larval stage, we have focused the remainder of our loss-of-function (LOF) studies using *form3-IR*.

### form3 promotes microtubule stabilization

Formins are well known regulators of the cytoskeleton and thus we hypothesized that *form3* functions in this process to modulate dendritic architecture. We utilized *CIV-GAL4* driven expression of transgenic multi-fluor cytoskeletal reporters in combination with *form3-IR* in order to simultaneously visualize the F-actin and MT cytoskeletons *in vivo*. These studies revealed disruptions in F-actin cytoskeletal distribution and a severe destabilization of dendritic MTs (Fig. 2A-B”). In contrast to the collapse of dendritic MT labeling, MT signal was still present on proximal CIV axons, albeit reduced relative to controls suggesting a more compartment-specific dendritic defect (Fig. 2A’,B’, arrowheads). To verify that the observed defects in MT stabilization were not due to a non-specific effect of *form3-IR* on expression of the mCherry tagged MAP Jupiter, we performed independent validation experiments. IHC analyses of CIV neurons expressing *form3-IR* were performed using antibodies against the *Drosophila* MAP1B molecule, Futsch. Compared to normal Futsch MT labeling in adjacent undisrupted CIII and CI da neurons in the same cluster, we observed a nearly complete loss of Futsch labeling in CIV dendrites expressing *form3-IR* (Fig. 2C-C”). These analyses corroborated the results of live imaging, collectively indicating that *form3* disruption leads to a collapse of MT architecture revealing a major functional requirement of Form3 in stabilizing dendritic MTs.

**Figure 2:**
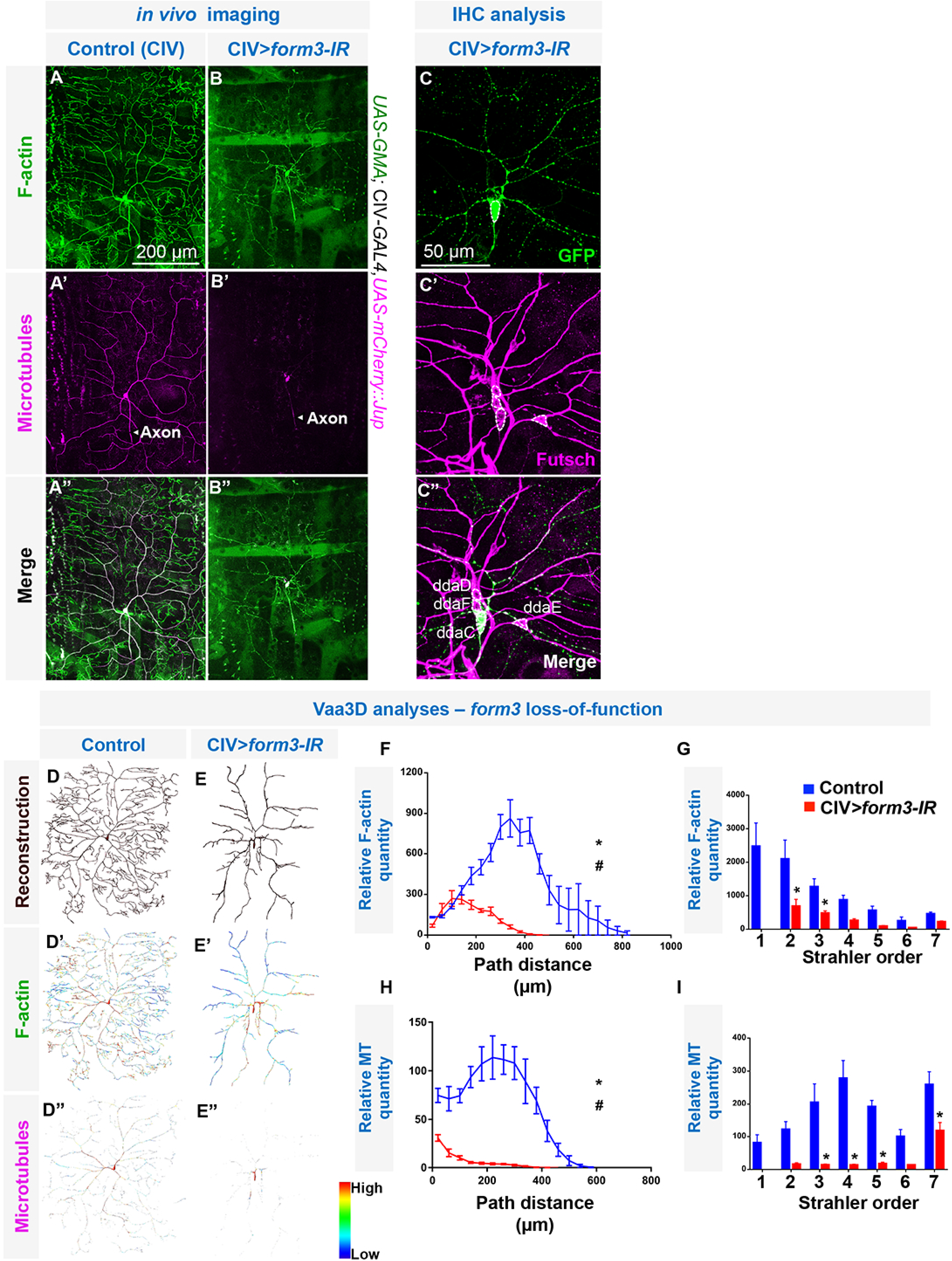
form3 functions in F-actin organization and microtubule stabilization. (**A-B”**) *CIV-GAL4* driven expression of *UAS-GMA* (which labels F-actin filaments via a GFP fused to the Moesin actin binding domain) and *UAS-mCherry::Jupiter* (which labels MTs via a mCherry fused to the MAP molecule Jupiter). (**A-A”**) Control CIV neuron. (**B-B”**) *form3-IR* CIV neuron reveals disruption of F-actin organization and dendritic MT stabilization. Representative images of N=10 CIV neurons per genotype. (**C-C”**) IHC of *GAL4^477^,UAS-mCD8::GFP >form3-IR* third instar larval filets double labeled with antibodies to the MAP1B protein Futsch to label MTs (**C’**) and GFP (**C**) to identify the affected dorsal cluster CIV neuron. The CI (ddaD/E) and CIII (ddaF) neurons exhibit normal, strong Futsch expression on MTs. Representative images of N=8 dorsal da neuron clusters. (**D-E**) Representative skeletons of the reconstructions of control and *form3-IR* CIV neurons, generated by a combination of Vaa3D, Neutube and TREES Toolbox. (D’,D”,E’,E”) Intensity maps of F-actin and MT generated in Vaa3D (N=5 for both control and *form3-IR)*. (**F,H**) Relative subcellular distributions of F-actin and MTs as a function of path distance from the soma; values are the mean (±SEM). (**G,I**) Relative F-actin quantity or relative MT quantity by Strahler order distribution, where 7=primary branch from cell body and 1=terminal branches; values are the mean (±SEM). Statistical tests performed in: (**F**) unpaired t-test for critical value (*) p=0.0191 and the corresponding radius (#) p=0.00005; (**H**) unpaired t-test for critical value (*) p=0.0060 and the corresponding radius (#) p=0.000443. (**G,I**) two-way ANOVA with Bonferroni correction: (**G_2_**) p=0.0035, (**G_3_**) p=0.0043; (**I_3,4,5_**) p=0.0001, (**I_7_**) p=0.0055.

To quantitatively assess the effect of *form3* disruption on the cytoskeleton, we employed newly developed next generation multi-channel neuronal reconstructions (Das et al. 2017). Multichannel reconstructions of cytoskeletal features revealed *form3-mediated* alterations in F-actin distribution and MT stabilization in LOF genetic backgrounds (Figure 2D-E”). Alterations in F-actin organization upon *form3* knockdown manifest as reduction in the critical value (the peak of the curve) and a proximal shift of the critical radius (the radius at which the peak of the curve lies) of the relative F-actin quantity (Figure 2F). We also found reductions in the relative F-actin quantity at several branch orders (Strahler order 2-5) with a complete absence of Strahler order 1 in *form3* knockdown, which makes up the majority of the F-actin rich terminal branches in control CIV neurons (Figure 2G). As predicted, the effect on MTs was severe as revealed by a collapse of MT architecture and overall reductions in the relative MT quantity as well as reductions at all branch orders (Figure 2H,I).

### form3 is required for complex dendritic arbor maintenance

To distinguish between potential roles in mediating dendritic specification vs. arbor maintenance, developmental time course studies were conducted examining CIV-specific *form3* knockdown at first, second and third instar larval stages. Time course analyses revealed that *form3-IR* disruption affects CIV dendritogenesis starting at early stages of development: however, the severity of the dendritic destabilization defect increases as the animal ages suggesting a major role in arbor maintenance (Figure 3A-H). Phenotypic comparisons at the first and second instar stages revealed an increased incidence of short interstitial branches emanating from the lower order dendrites in *form3-IR* relative to control (Figure 3A,A’B,B’,E,E’,F,F’) whereas third instar *form3-IR* neurons exhibited dramatic reductions in dendritic branches (Figure 3C,C’,G,G’). These qualitative observations are confirmed by morphometric analyses of *form3-IR* which revealed that while there is only a modest or no difference in the number of branches at first and second instars stages when compared to control, there is severe reduction in the number of branches at the third instar stage (Figure 3I). Similarly, branch density is most severely reduced at the third instar stage (Figure 3J). Collectively, these data are indicative of a progressive degeneration of terminal branches over the developmental time course (Figure 3D,H).

**Figure 3:**
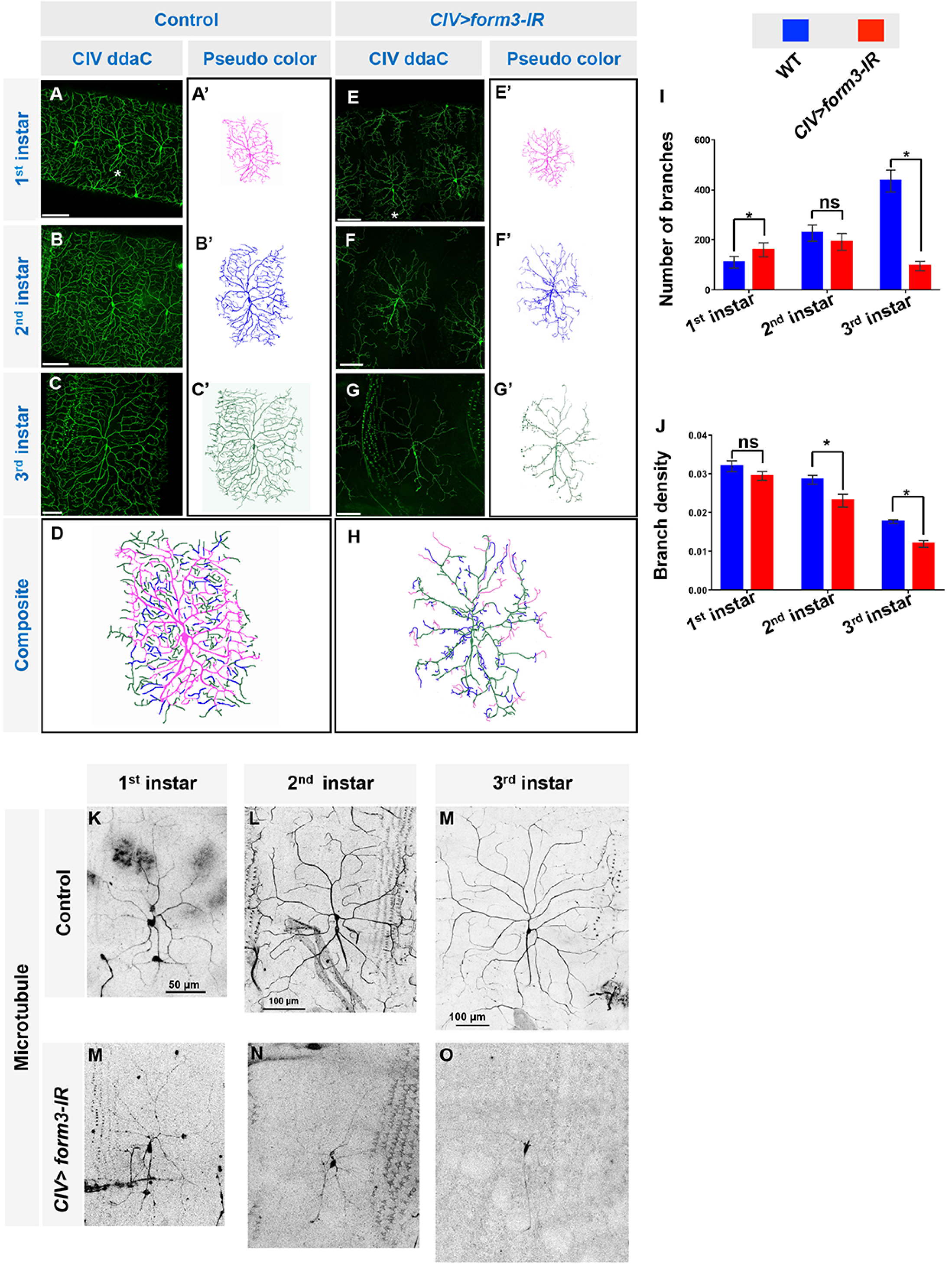
form3 is required for dendritic arbor maintenance. (**A-H**) Representative larval developmental time course images of the same CIV neuron taken of control (**A,B,C**) and *form3-IR* (**E,F,G**) at 1^st^ instar, 2^nd^ instar and 3^rd^ larval instar stages, respectively. (A’-C’) Pseudo-colored panels show the same neuron color coded for their developmental stages (magenta as 1^st^, blue as 2^nd^ and green as 3^rd^ instar). (**D**) Composite image shows the arborization of the dendritic branches that occurred from 1^st^ to 2^nd^ to 3^rd^ instar stages, where blue marks those dendrites added from 1^st^ to 2^nd^ instars, and green marks those dendrites that were added from 2^nd^ to 3^rd^ instars. (**H**) Composite image shows the loss of the dendritic branches that occurred from 1^st^ to 2^nd^ to 3^rd^ instar stages, where magenta marks those dendrites that were lost from 1^st^ to 3^rd^ instar, and blue marks those dendrites lost from 2^nd^ to 3^rd^ instars. Note, in the panel D and H the branches are not exact to scale. (**I,J**) Quantitative analyses measuring number of branches and branch density of control (N=7) and *form3-IR* (N=7) at different stages of the developmental time course as indicated on the figure. (**K-O**) *CIV-GAL4* driven expression of *UAS-mCherry::Jupiter* in control (N=9) (**K-M**) or *form3-IR* (N=8) (**M-O**) at different stages of the developmental time course as indicated on the figure. Statistical tests performed in: (**I,J**) unpaired t-test, (I_1st-instar_) p=0.004164 and (Winstar) p=4.98481E-08; (J_2nd-instar_) p=0.022424 and (J_3rd-instar_) p=0.000246. NS represents not significant p>0.05. Data are represented as mean ± SEM.

Given the effect of *form3-IR* on MT stability at the third instar stage and the progressive loss of terminal branches observed in time course analyses, we hypothesized that loss of dendritic branches would correlate with progressive destabilization of MTs over development. Consistent with this hypothesis, we find that MT destabilization is likewise progressive over the developmental time course from first to third instar larval stages (Figure 3K-O). These results indicate that progressive MT destabilization strongly correlates with loss of higher order dendritic branching.

### form3 regulates higher order branching

As *form3* disruption produces severe defects in higher order dendritic branching, we sought to test a hypothesis that *form3* overexpression will lead to excessive terminal branching. We overexpressed *UAS-form3* in CIV, which revealed a shift in branching distribution characterized by a reduction in branching proximal to the cell body as well as exuberant terminal branching and elongated terminal dendrite extension (Fig. 4A-B’). Sholl analysis revealed a significant increase in the critical value proximal to the termini compared to controls (Figure 4C), which occurs prominently around 105-200 micron from the terminals (Figure 4D). Moreover, Strahler analysis revealed a significant increase in number of terminal branches indicated by Strahler order 1 (Figure 4E).

**Figure 4:**
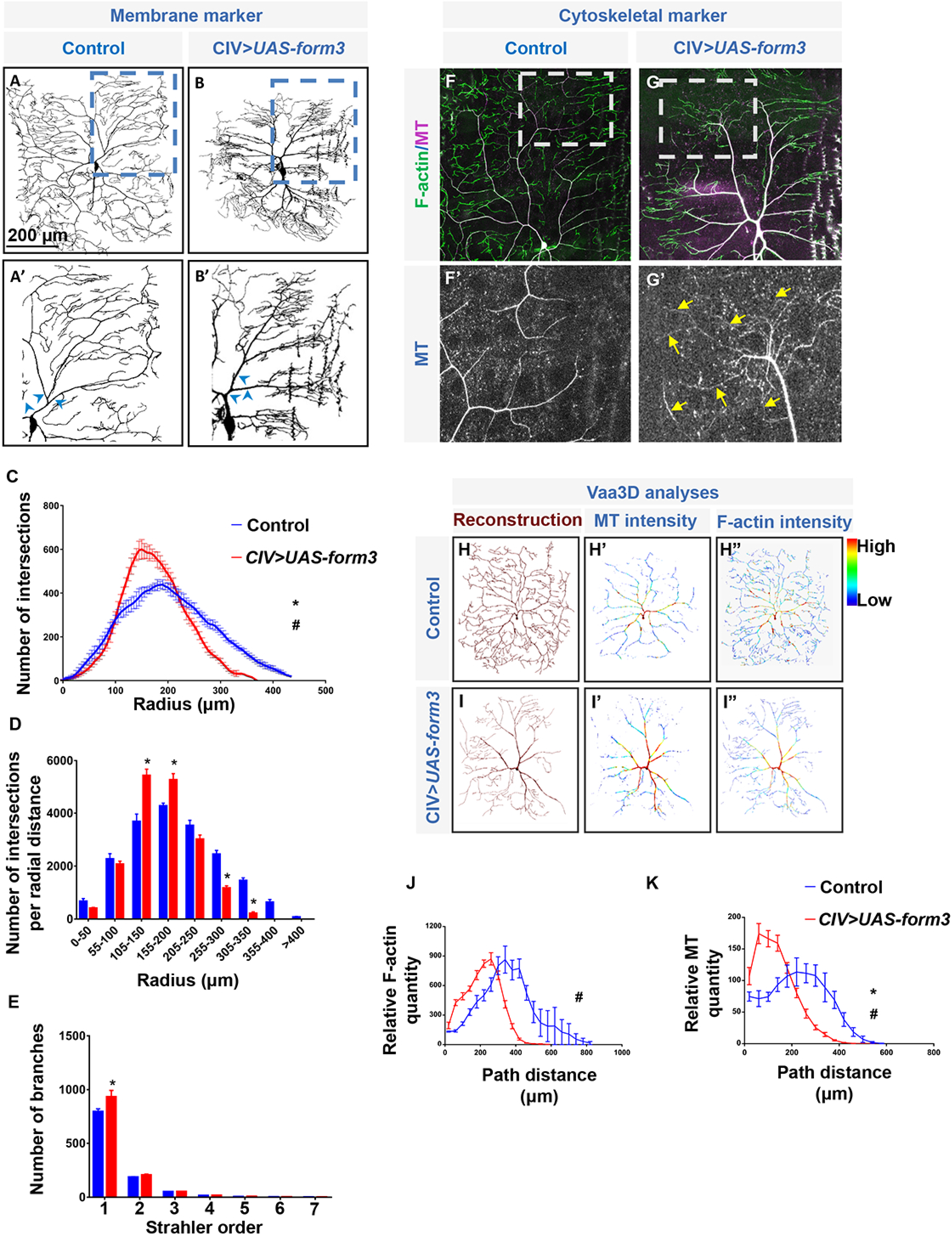
Form3 overexpression promotes excessive terminal branching and elongation by promoting or stabilizing MT extension into these processes. (**A,A’**) Representative image of control CIV neurons (N=12). (**B,B’**) Representative image of *form3* overexpression CIV neurons (N=7). Relative to control CIV neurons (**A,A’**), *form3* overexpression (**B,B’**) leads to a distal shift in branching complexity characterized by excessive growth of terminal dendritic branches and thickened lower order (e.g. primary, secondary branches, indicated by blue arrowheads). (**C,D**) Sholl profile of control (N=12) vs. *form3* overexpression (N=7). Values are the mean (±SEM) for the number of intersections as a function of radius distance from the termini (zero) to the cell body of the neurons. (**D**) Sholl profiles for control vs. *form3* overexpression data plotted as histogram to reflect the total number of intersections per corresponding radial distance (Euclidean) from the termini (zero) to the soma highlighting local effects on branch distributions. (**E**) Reverse Strahler analysis of control vs. *form3* overexpression. Values are the mean (±SEM) for the number of dendritic branches in each branch order (Strahler order), where 7=primary branch from cell body and 1=terminal branches. (**F,F’**) Dendritic terminals of control CIV neurons (N=10) rich in F-actin processes. (**G,G’**) Dendritic terminals of CIV neurons overexpressing Form3 (N=10) are populated by F-actin processes, but also display increased MT invasion into these processes. (**H-I”**) Multichannel reconstructions of cytoskeletal features in control (N=5) and *form3* overexpression (N=8) CIV neurons. Heat maps represent the relative distribution and intensity of F-actin and MTs in controls vs. *form3* overexpression. Note that *form3* overexpression leads to increased MT intensity and abnormally increased diameters of primary dendritic branches emerging from the cell body. (**J,K**) Relative subcellular distributions of F-actin and MTs as a function of path distance from the soma. Statistical tests performed in: (**C**) unpaired t-test for critical value (*) p=0.00825 and the corresponding radius (#) p=0.00041; (**D, E**) two-way ANOVA with Bonferroni correction: (**D_105-150_**) p= 0.0001, (**D_155-200_**) p=0.0005; (**D_305-350_**) p=0.0001, (**E_1_**) p=0.0001; (**J**) unpaired t-test for corresponding radius (#) p=0.002528; (**K**) unpaired t-test for critical value (*) p=0.024318 and for corresponding radius (#) p=0.000419. Data are represented as mean ± SEM.

Another striking feature observed with *form3* overexpression is a change in primary dendrite branch thickness. Relative to control, primary branch diameter was notably increased by *form3* overexpression (Figure 4A’B’, arrowheads), suggesting increased Form3 levels may alter underlying cytoskeletal organization. Therefore, we sought to assess the effect of *form3* overexpression on the cytoskeleton. Multichannel cytoskeletal reconstructions revealed an altered F-actin signature in CIV neurons overexpression Form3, which when quantified showed a leftward shift of the critical value (peak of the curve) indicating an increase in the F-actin in the first few dendritic branch orders (Figure 4H”,I”,J). Furthermore, cytoskeletal analyses revealed Form3 overexpression leads to an increase in the MT intensity profile proximal to the cell body (Figure 4F,G,H’,I’) indicative of a role of Form3 in promoting MT density in primary dendritic branches. This raises the question of whether there may be changes in the cytoskeletal organization of dendritic termini as well. Relative to control termini (Figure 4F,F’), Form3 overexpression analyses revealed increased MT invasion into the elongated terminal branches likely stabilizing their extension (Figure 4G,G’).

### form3 directly interacts with microtubules and promotes microtubule acetylation

Based on the regulatory relationship between Form3 function and MT architecture, we sought to examine whether this may occur via direct or indirect mechanisms. To address this, we performed MT co-sedimentation (±MTs) and MT precipitation assays using GST-tagged Form3 constructs expressing either the FH1-FH2 domains or the FH2 domain alone as compared to GST controls lacking Form3 sequences. Analyses of FH1-FH2 fusion proteins revealed specific co-sedimentation in pellet fractions only in the presence of MTs (Figure 5A) indicative of a direct interaction. Analyses of the FH2 domain alone revealed low level self-pelleting in the absence of MTs, however despite this, there was a significant increase in the amount of FH2 fusion protein that co-sedimented with MTs specifically indicating the FH2 domain is sufficient for Form3-MT direct interaction. To further validate these results, we performed MT precipitation assays revealing tubulin precipitated only in the presence of the Form3 constructs, but not with GST alone (Figure 5B).

**Figure 5:**
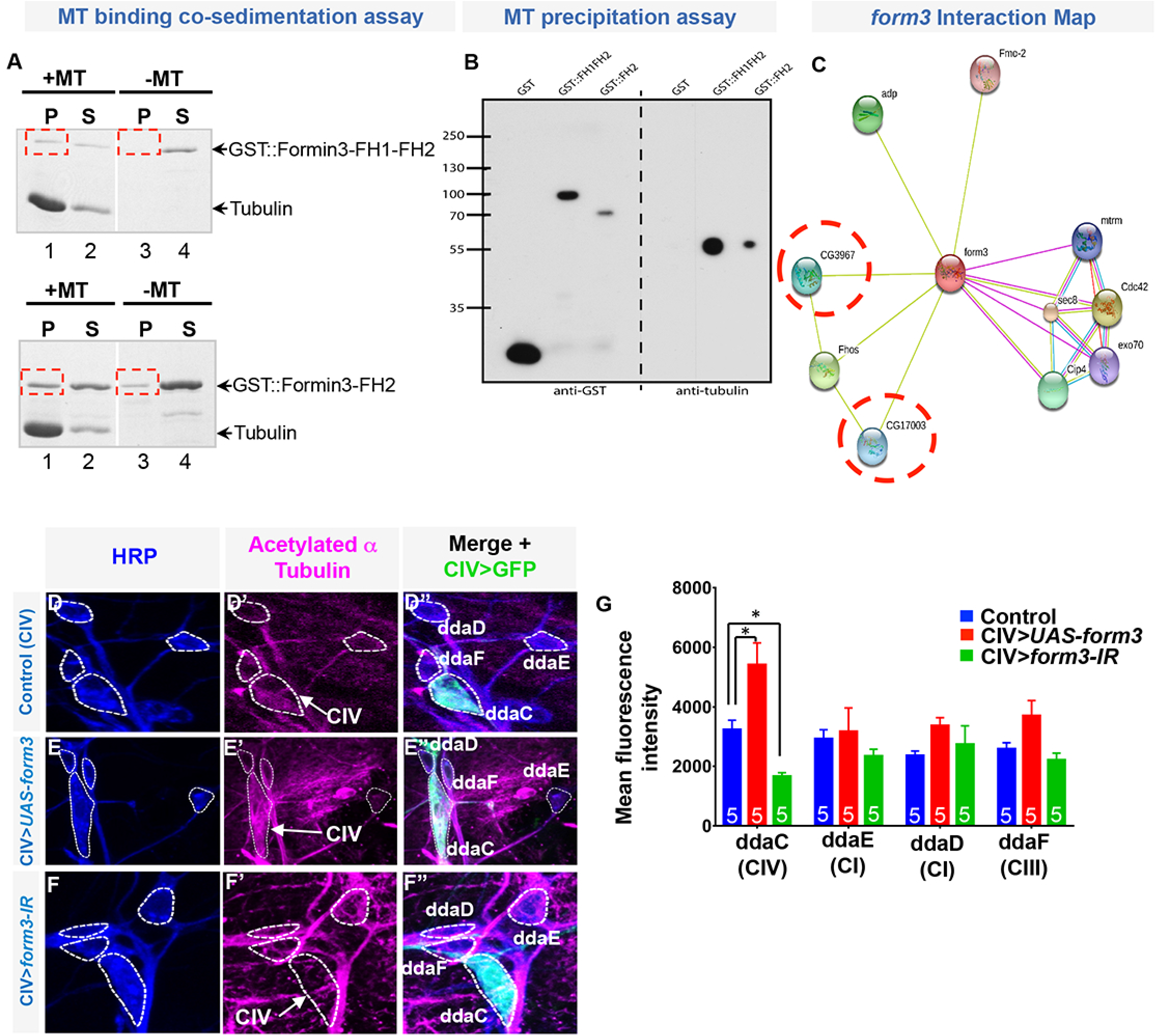
form3 directly interacts with microtubules and promotes microtubule acetylation. (**A**) MT co-sedimentation assay. GST-tagged Form3 constructs expressing either the FH1-FH2 domains or the FH2 domain alone incubated in the presence or absence of MTs. Western blots probed for anti-GST and anti-α-Tubulin. The FH1-FH2 fusion displays specific co-sedimentation in pellet fractions only in the presence of MTs, whereas the FH2 fusion displays a low level of self-pelleting that is notably increased in the presence of MTs. Pellet fractions indicated by dashed red boxes, P=pellet, S=supernatant. (**B**) MT precipitation assay. Western blots probed for anti-GST or anti-α-Tubulin. Tubulin co-precipitates only in the presence of the Form3 GST fusion constructs, but not with the GST construct alone. Protein molecular weights indicated at left in kDa. Biochemical assays performed in duplicate. (**C**) Interaction map for predicted/knownform3-interacting genes. *CG17003* and *CG3967* in red circles belong to *ATAT1* family. Interaction map data obtained from <u>http://string-db.org/</u>. (**D-F”**) Third instar larval filets triple labeled for HRP to mark all da sensory neurons (**D,E,F**); anti-acetylated α tubulin (**D’,E’,F’**) and anti-GFP to mark CIV neurons (**D”,E”,F”**). Relative to controls (N=5) (**D-D”,G**), CIV *form3-IR* (N=5) knockdown leads to a reduction in acetylated α tubulin (**F-F”,G**), whereas *form3* overexpression (N=5) results in a strong increase in acetylated α tubulin (**E-E”,G**). Statistical test performed: (**G**) two-way ANOVA with Bonferroni correction (**G_ddaC_**) p=0.002 (CIV>*UAS-form3*) and p=0.032 (CIV>*form3-IR*). Data are represented as mean ± SEM.

Previous studies demonstrate the ability of Formin proteins to induce MT acetylation (Thurston et al. 2012), however whether this is true for Form3 remains an open question. Interestingly, STRING analyses revealed predicted *form3* interactions with the two *Drosophila* α-tubulin N-acetyltransferase (*ATAT1*) family members, *CG3967* and *CG17003* (Figure 5C), and CIV-specific microarray analyses revealed that both of these molecules are significantly expressed in these neurons (data not shown; Iyer et al. 2013). This suggests the possibility that Form3 may stabilize MTs by interacting with ATAT1 proteins to promote acetylation, and thereby stabilization, of dendritic MTs.

To characterize the putative *form3* interactors predicted by STRING, we performed CIV-specific knockdown analyses. Select interactors appear to have a role in CIV branching and terminal dendritic patterning, including *exo70* and *Sec8*, both linked to exocyst complex, whereas disruptions in *mtrm* and the Formin *Fhos* had mild to no effect on CIV dendritogenesis (Figure. 5-1B,C,F-H; Figure 1-1D,G,H). Previous studies have implicated *Rop* together with other components of the exocyst complex (*Sec5, Sec6*) in promoting dendritic growth and maintenance (Peng et al. 2015). Knockdown of *CG17003* or *CG3967* had modest, but insignificant effects on dendritic branching and growth (Figure 5-1D,E,G,H). There is, however, a formal possibility that these two ATAT1 molecules exhibit functional redundancy in MT acetylation. Nevertheless, these findings, coupled with previous evidence in other systems (Thurston et al. 2012), prompted us to directly investigate how *form3* disruptions may impact post-translational modifications that could contribute to MT de/stabilization.

**Extended Figure 5-1:**
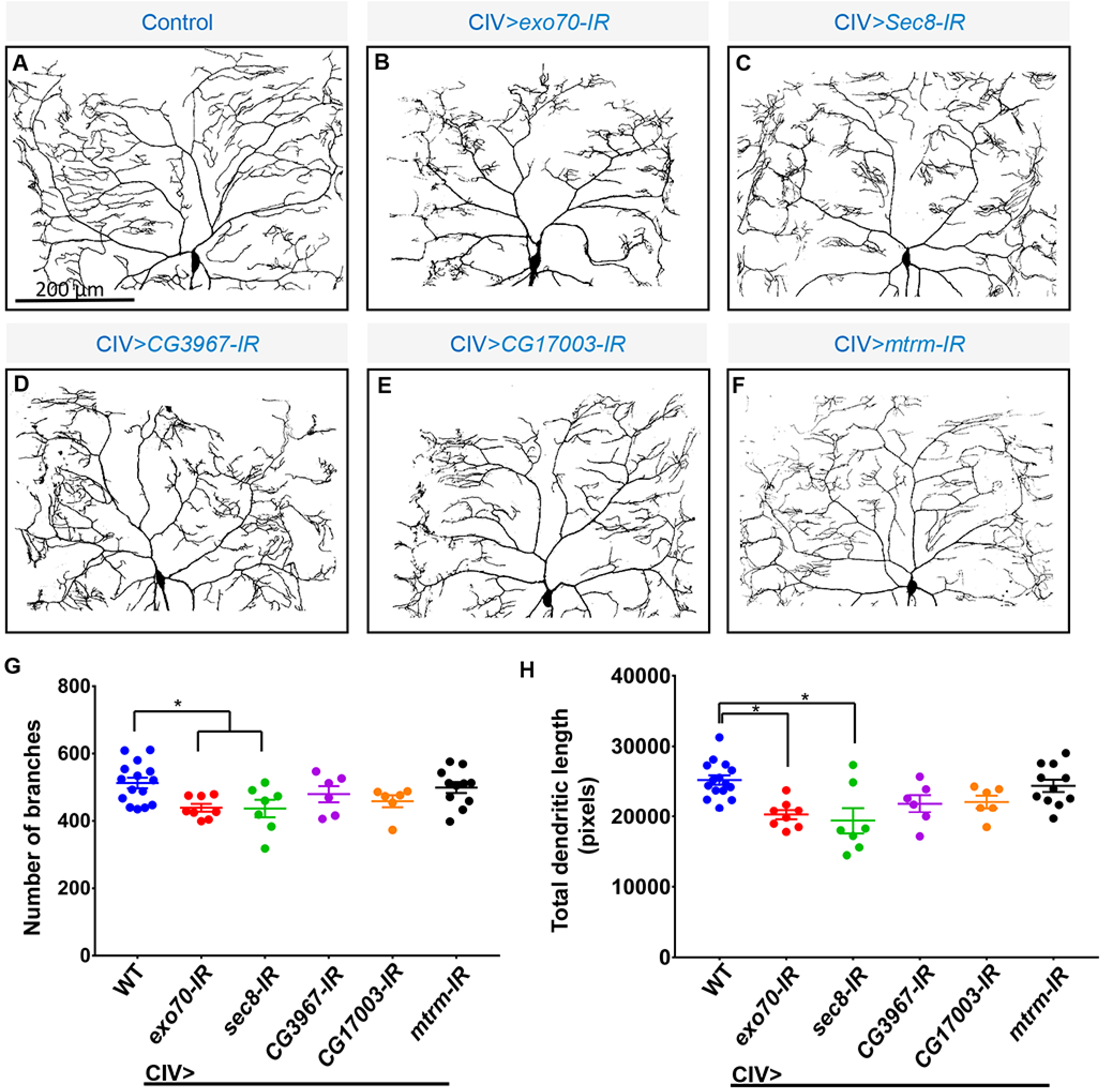
Phenotypic analyses of putative form3 interactors. (**A-F**) Representative images of CIV dendrites in control (N=15) (**A**) and (**B-F**) RNAi knockdowns for putative *form3* interactors: *exo70* (N=8); *Sec8* (N=7); *CG3967* (N=6); *CG17003* (N=6); *mtrm* (N=11). (**G,H**) Quantitative analyses. Statistical test performed: (**G,H**) One-way ANOVA with Bonferroni correction; (**G**) p=0.0193 *(exo70-IR)*, p=0.0220 *(Sec8-IR);* (**H**) p=0.002 (*exo70-IR*), p=0.0005 (*Sec8-IR*).

To investigate this, we performed IHC analyses in *form3* LOF and overexpression genetic backgrounds. The results demonstrated that CIV-specific *form3* knockdown leads to a reduction of acetylated α-tubulin, while overexpression increases the levels of acetylated α-tubulin (Fig. 5D-G). Increased levels of acetylated tubulin may explain, in part, why Form3 overexpression leads to excessive terminal branching and elongation, as well as thickening of proximal branches, whereas reductions in acetylated tubulin may explain, in part, why *form3* mutant neurons exhibit MT destabilization.

### form3 is required for dendritic organelle trafficking and architecture

Collective evidence implicates Form3 in MT stabilization, which while clearly critical to supporting dendritic complexity does not fully explain why the arbor exhibits progressive degeneration over developmental time. Therefore, we sought to examine the potential functional consequences of a destabilized MT cytoskeleton by examining trafficking of organelles essential for supporting dendritic development. Studies have demonstrated that defects in mitochondrial trafficking can lead to dendritic degeneration/fragmentation in both invertebrates and vertebrates (Tsubouchi et al. 2009; Lopez-Domenech et al. 2016). Moreover, INF2 (human ortholog of Form3) has been shown to affect mitochondrial length and ER-mitochondrial interactions (Korobova et al. 2013), thus Form3 may function in mitochondrial dynamics. *In vivo* imaging revealed specific inhibition of mitochondria dendritic trafficking in *form3-IR* whereas proximal axonal trafficking appears largely normal (Fig. 6A-B”). We also observed a significant decrease in the dendritic mitochondria index revealing that there are also fewer dendritic mitochondria when normalized to total dendritic length (Figure 6C). These data indicate that mitochondrial trafficking dynamics is in part dependent on Form3, as well as a stable MT cytoskeleton.

**Figure 6:**
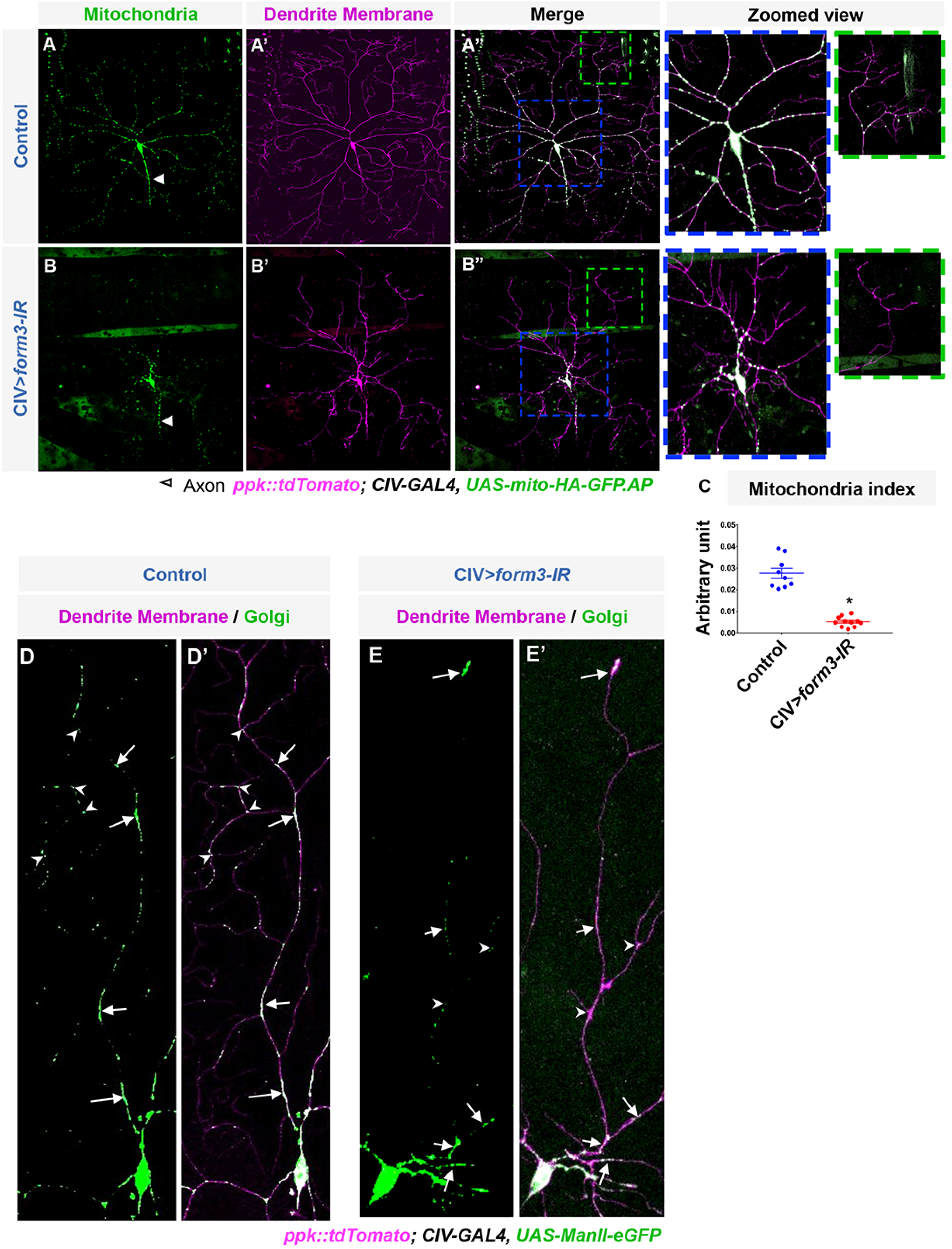
form3 is required for dendritic organelle trafficking. Relative to controls (N=9) (**A-A”**), *form3-IR* (N=11) CIV neurons (**B-B”**) exhibit dendritic collapse (**B’**) and a severe inhibition of mitochondria trafficking onto dendrites (**B,B”**), but not axons (arrowheads). Zoomed view of the corresponding colored boxes, where blue box represents the area proximal to the cell body and green box represents the area distal to the cell body. (**C**) Mitochondria index: number of dendritic mitochondria divided by total dendritic length in pixels (mean ± SEM). Relative to controls (N=5) (**D-D”**), *form3-IR* (N=6) CIV neurons (**E-E”**) exhibit defects in dendritic Golgi trafficking and aberrant Golgi architecture suggesting fragmentation (**E’**). (**D-E’**) Arrows denote locations of interstitial “islands” of fused Golgi (control) or fragmented Golgi puncta *(form3-IR*); arrowheads denote dendritic branch points. Statistical test performed: (**C**) unpaired t-test, p=2.77142E-05.

Trafficking of satellite Golgi has been demonstrated to play key roles in regulating dendritic development and MT nucleation (Ye et al. 2007; Ori-McKenney et al. 2012). We hypothesized that *form3* disruption may impair proper trafficking of satellite Golgi due to a destabilized MT cytoskeleton, which could, in part, contribute to the observed dendritic retraction. We discovered aberrant trafficking in *form3-IR* neurons with the majority of the satellite Golgi confined to the proximal branches though we did observe some distal Golgi trafficking to termini. The overall number of satellite Golgi was notably reduced when two analogous branches of control and *form3-IR* were compared (Figure 6D-E’). In control, there was a distinct localization of satellite Golgi to dendritic branch points, which was often not the case in *form3-IR*. In addition, while controls display “islands” of fused Golgi located along interstitial branches, such islands are not observed with *form3-IR*, but rather only small puncta, indicative of Golgi fragmentation (Figure 6D,E). These data suggest that an intact MT cytoskeleton is required for normal satellite Golgi translocation, and that defects in trafficking may contribute to overall dendritic atrophy observed in *form3-IR* neurons.

### INF2 can functionally substitute for form3 in promoting dendrite complexity and MT stability

To investigate the hypothesis that *Drosophila* Form3 and human INF2 may share conserved functions, we generated transgenic fly strains to allow for inducible expression of either full length INF2 or a truncated version in which autoinhibitory regulatory domains (DID/DAD) were deleted, leaving only the FH1 and FH2 domains (FH1-FH2) (Figure 7A). Previous studies in *C. elegans* revealed that removal of the DID/DAD inhibitory domains was required for INF2 rescue of the worm ortholog *exc-6* (Shaye and Greenwald 2015) and Form3 protein does not contain the DID/DAD domains.

**Figure 7:**
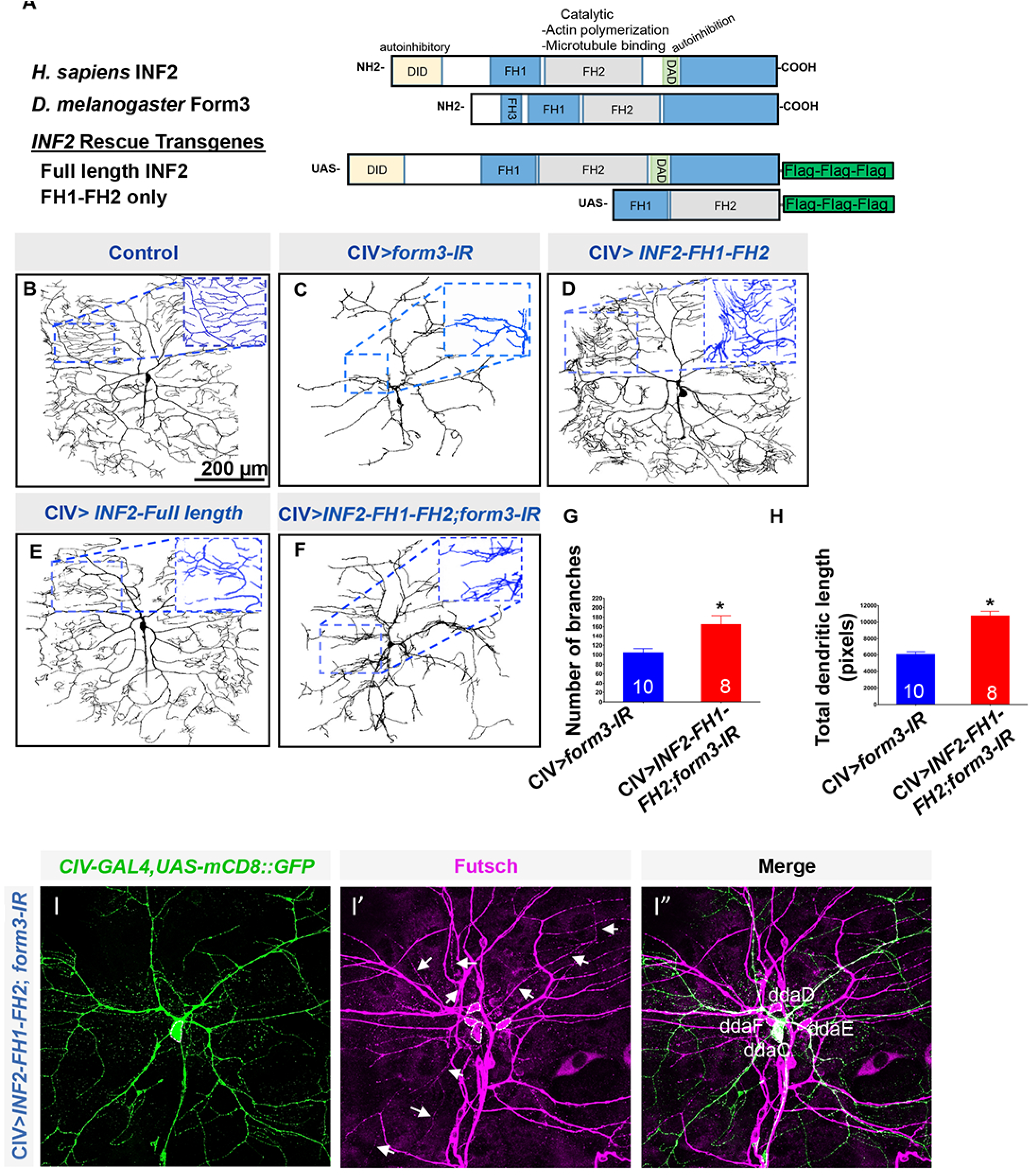
INF2 can functionally substitute for form3 in promoting dendrite complexity and MT stability. (**A**) Schematic diagrams of human INF2 and *Drosophila* Form3 domain organization; schematic diagrams of *INF2* rescue transgenes. (**B**) Control CIV neurons (N=10). (**C-F**) CIV-specific expression of the corresponding transgenes (N=10). (**F**) CIV-specific expression of *form3-IR* with simultaneous expression of *INF2-FH1-FH2* in the same neuron (N=8). (**G,H**) Quantitative rescue measurements of number of branches and total dendritic length (mean ± SEM). (I-I”) CIV-specific expression of *INF2-FH1-FH2* in *form3-IR* background (N=8). Introduction of *INF2-FH1-FH2* rescues MT signal labeled with anti-Futsch (magenta). Arrows in panel I’ depict rescued labeling of CIV dendrite MTs. Statistical tests performed: (**G**) unpaired t-test p=0.0001; (**H**) unpaired t-test p=0.0002.

We initiated our *INF2* analyses by first examining the functional consequences of overexpressing these transgenes in CIV neurons. Overexpression of the truncated *INF2-FH1-FH2* transgene induced changes in dendritic architecture that are phenotypically consistent with *form3* GOF. In both Form3 and INF2-FH1-FH2 overexpression, we observed a shift in branch distribution towards dendritic terminals, which are abnormally hyperproliferated and elongated suggesting similar functional effects (Figures, 4B,B’ and 7D). In contrast, full length INF2 overexpression had no significant effects on CIV dendritic morphology, likely due to autoinhibition (Figure 7B,E).

To determine whether INF2 can functionally substitute and rescue *form3-IR* defects in CIV dendritogenesis we focused rescue analyses on the *INF2-FH1-FH2* transgene given the results from the overexpression studies and previous rescue studies of the worm ortholog *exc-6*. The *INF2-FH1-FH2* variant was introduced into the *form3-IR* genetic background and out-crossed to a *CIV-GAL4* driver. These analyses revealed a partial rescue of CIV morphological defects (Figure 7C,F) partially reverting lost dendritic complexity by promoting increased dendritic growth and branching (Fig. 7C,F,G,H).

As *form3* defects lead to dendritic MT destabilization (Figure 2), we hypothesized that introduction of *INF2-FH1-FH2* would potentially rescue this defect. In contrast to *form3-IR*, introduction of *INF2-FH1-FH2* in the *form3-IR* background resulted in a rescue of dendritic MTs (Figure 7I-I”). These studies demonstrate that INF2-FH1-FH2 is not only capable of partially rescuing the overall dendritic complexity defects, but can also recover dendritic MT stability both proximal and distal to the cell body (Figure 7J’, arrows). Collectively, these analyses support conserved functional roles for Form3 and INF2, via FH1-FH2, in promoting dendritic complexity and stabilizing MTs.

### Roles of form3 and INF2 in peripheral sensitivity and thermosensory nociception

CIV neurons function as polymodal nociceptors and are required to mediate a variety of nocifensive responses including an aversive body rolling (360°C) behavior in larvae upon exposure to noxious heat or mechanical stimuli (Tracey et al. 2003; Himmel et al. 2017). Furthermore, recent evidence connects defects in dendritic architecture with noxious heat-evoked nocifensive behavioral sensitivity (Honjo et al. 2016). We hypothesized that *form3* disruption in these neurons may lead to reduced peripheral sensitivity to noxious heat and thereby impair nociceptive behavior. Thus, we expressed *form3-IR* in CIV neurons and observed nociceptive behavioral responses when challenged with a noxious heat (45°C) stimulus. These analyses revealed a severe impairment in noxious heat evoked behavioral responses (Figure 8C,E,F; Movie 1), thereby revealing a dramatic loss of peripheral sensitivity. This behavioral defect was not due to any general defect in locomotion as both control and *form3-IR* larvae exhibit normal locomotor behavior. In contrast to control larvae, CIV-specific inhibition of *form3* function leads to a dramatic increase in the latency to respond (among those few larvae which ever respond) (Figure 8A,C,E). The majority of *form3-IR* larvae were classified as non-responders as they fail to exhibit rolling or other documented nociceptive behaviors within the 20 second assay period (Figure 8C,F).

**Figure 8:**
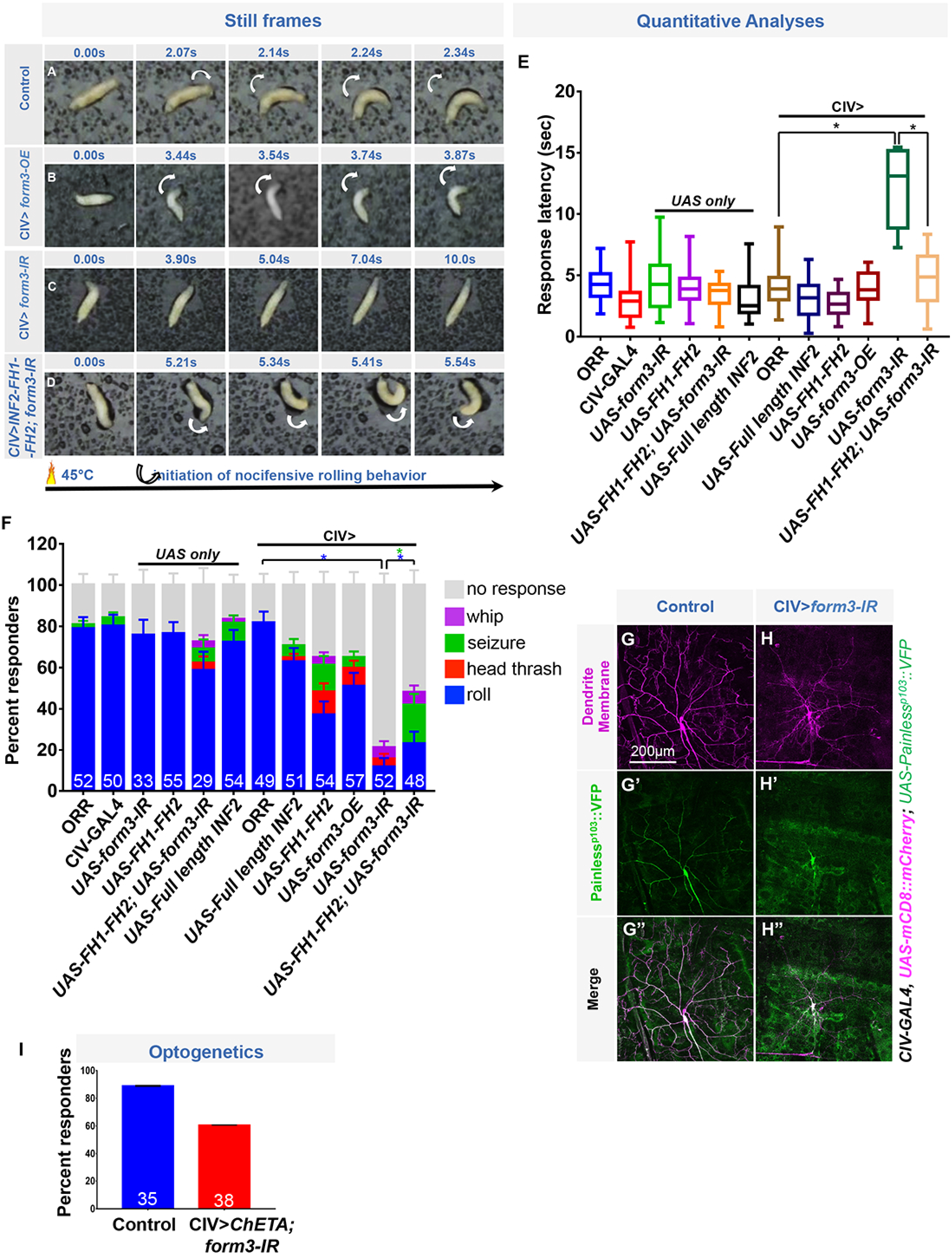
form3 is required for noxious heat nociception, which can be rescued by introduction of INF2-FH1-FH2. (**A-D**) Representative still images of noxious heat-evoked rolling behavior for the designated genotypes. (**E**) Latency to roll in seconds (mean ± SEM) for the designated genotypes at 45°C. (**F**) Percent responders (mean ± SEP) for the respective nocifensive behaviotypes in the designated genotypes at 45°C. Number of third instar larvae (**N**) for (**E,F**) by genotype is represented on the bar graph displayed in (**F**). (G-H”) Representative images of the CIV-specific expression of *UAS-painless^p1Ω3^::VFP* in control (N=10) or *form3-IR* (N=7) neurons, where dendrite membrane is labeled with *UAS-mCD8::mCherry* in magenta (**G,H**) and Painless^p103^::VFP in green (G’,H’). (**I**) Percent responders (mean ± SEP) in CIV-specific optogenetic activation. Statistical test performed: (**E**) one-way ANOVA with Bonferroni correction, comparisons are indicated on the figure. (**E**) p=0.0001 (CIV>*UAS-form3-IR*) and p=0.0001 (CIV>*UAS-FH1-FH2; UAS-form3*). (**F**) A two sample t-test between proportions p=0.0001 (CIV>*UAS-form3-IR*) and p=0.00001 (CIV>*UAS-FH1-FH2; UAS-form3-IR* for roll) and p=0.00001 (CIV>*UAS-FH1-FH2; UAS-form3-IR* for seizure).

Next, we sought to determine whether *form3* or *INF2-FH1-FH2* overexpression may lead to changes in nociceptive behavior when challenged with noxious heat. We found that neither *form3* nor *INF2-FH1-FH2* overexpression resulted in a significant change from controls with respect to latency (Figure 8B,E); however, we did observe a reduction in the percentage of responders in both these conditions (Figure 8F). Apart from nocifensive rolling, several other noxious heat-evoked behaviotypes have been described, including larval whipping, head thrashing, and seizures (Chattopadhyay et al. 2012). Relative to controls, and among those larvae that failed to execute rolling behavior, these other behaviors were variably observed with greater incidence in *form3* and *INF2-FH1-FH2* overexpression, as well as to some modest extent in *form-IR* (Figure 8F). This may suggest that the alterations in dendritic morphology observed with *form3* or *INF2-FH1-FH2* manipulations may impair normal processing of thermal stimuli resulting in aberrant behavioral repertoires.

Intriguingly, mutations in *INF2* have been causally linked to CMT disease, though the mechanisms of action in CMT pathology are incompletely understood (Boyer et al. 2011). Neurological features of CMT include peripheral motor and sensory neuropathies, and the primary phenotypes consist of progressive distal muscle weakness and atrophy, reduced tendon reflexes, foot and hand deformities and peripheral insensitivity (Ekins et al. 2015). CMT sensory neuropathies lead to distal sensory loss resulting in a reduced ability to sense heat, cold, and pain, yet the neural bases of these sensory defects remains to be fully elucidated. Given the causative role of *INF2* mutations in CMT disease and the impaired distal sensation to thermal stimuli in CMT patients, we next tested the hypothesis that introduction of the *INF2-FH1-FH2* transgene into the *form3-IR* background may rescue the impaired behavioral responses to thermal nociceptive stimuli. Consistent with our hypothesis, we found that CIV expression of *INF2-FH1-FH2* significantly rescued the behavioral latency defects observed in *form3-IR* larvae leading to an increase in the percentage of behavioral responders (Figure 8D-F; Movie 2). While the introduction of INF2-FH1-FH2 only partially rescues the behavioral defects, it nonetheless suggests a conserved role for Form3 and INF2 in regulating peripheral sensitivity to nociceptive stimuli that has potential implications for the etiological bases of CMT peripheral sensory neuropathy.

Previous studies have linked the TRPV4 channel in human to CMT2C due to disruptions in its Ankyrin repeats (ARs) (Landouré et al. 2010). Noxious heat detection in *Drosophila* larvae requires the function of two TRPA channels, Painless and TRPA1 both of which contain Ankyrin repeats (ARs) (Tracey et al. 2003; Zhong et al. 2012). In the mechanosensing TRP channel NompC, the ARs provide direct linkage of the channel to the MT cytoskeleton and are required for mechanical gating of the channel (Zhang et al. 2015). Moreover, studies of the canonical Painless protein which bears N-terminal ARs have revealed that these domains are necessary for thermal, but not mechanical, nociception (Hwang et al. 2012). These findings led us to hypothesize that *form3-mediated* defects in the MT cytoskeleton may affect trafficking or localization of AR containing TRP channels such as Painless. To investigate this hypothesis, we conducted *in vivo* imaging of CIV neurons expressing *UAS-Painless^p103^::VFP* while simultaneously expressing *form3-IR*. In contrast to controls in which Painless^p103^::VFP is strongly expressed on CIV cell bodies, as well as throughout dendrites and proximal axons (Figure 8G-G”), in *form3-IR* neurons there is a strong inhibition of Painless trafficking onto dendrites distal to the cell body despite comparable levels of cell body expression and to a lesser extent axonal expression (Figure 8H-H”). These data suggest that form3-mediated defects in nociceptive behavior may be due, in part, to aberrant trafficking of noxious thermosensory channels such as Painless and that Painless dendritic trafficking is at least somewhat dependent on a stable MT cytoskeleton.

Finally, to assess the putative functional role(s) of Form3 in noxious heat-evoked behavior, we examined whether Form3 is required for the sensory transduction of noxious thermal stimuli or action potential (AP) propagation via optogenetic activation studies. If Form3 is required at the sensory transduction stage, then optogenetic activation of CIV neurons should largely bypass the *form3-IR* defect and evoke the stereotypical rolling response, whereas if Form3 functions in AP propagation, then optogenetic activation will be insufficient to elicit the rolling behavior. These analyses revealed ~60% of the *form3-IR* larvae performed rolling behavior with optogenetic activation (Figure 8I; Movie 3). While we observed that *form3-IR* larvae had a somewhat longer latency to roll upon blue light exposure, and not all larvae exhibit a roll, this is likely attributable to the severe reduction in dendritic field coverage, which has been shown to impair nociceptive behavior (Honjo et al. 2015). These data support a primary role for *form3* in sensory transduction rather than AP propagation.

## Discussion

### form3 function in dendrites and MT stabilization

LOF analyses revealed *form3* is required for higher order branching complexity in CIV nociceptive sensory neurons; however lack of gross defects in CIV axon patterning suggests a specific role of Form3 in stabilizing CIV dendrites. This raises interesting questions regarding the dendritic specificity of *form3* function. While we do not yet fully understand this compartment specificity in terms of *form3* function, we speculate that Form3 perhaps interacts with different partners in these dendritic vs. axonal compartments, ultimately impacting its overall function.

Dendritic development is a complex phenomenon, which requires spatio-temporal regulation of local cytoskeletal interactors to direct specific morphological features of the neuron. Molecules involved in this process can have one of many roles, such as arbor specification, growth by enhancement, suppression by reduction or simply maintenance of the dendritic arbor. We demonstrate that Form3 is required in dendritic arbor maintenance as *form3* mutants exhibit progressive dendritic atrophy ultimately leading to a highly rudimentary arbor and dramatically reduced dendritic field coverage.

The hypothesis that Formins may regulate MTs has been proposed for some time, however, only recently have biochemical studies begun to explore how Formins interact with MTs and affect their dynamic properties. Studies have shown that multiple mammalian Formins, as well as the fly Formin, Capuccino, can interact directly with MTs (Breitsprecher and Goode 2013; Roth-Johnson et al. 2014); however whether this is a specific or more general property of Formin molecules remains unclear. At a mechanistic level, multiple converging lines of evidence implicate Form3 in primarily regulating MT stabilization including direct interaction with MTs and promotion of MT acetylation. Consistent with this, previous studies demonstrate that formation of stabilized MTs requires INF2 (Andrés-Delgado, 2012).

### form3 function in dendritic organelle trafficking and architecture

Proper mitochondrial function is required to support neuronal development and function as mitochondria are fundamentally important for several cellular events, such as ATP production, Ca^2+^ regulation, as well as release and reuptake of neurotransmitters at synapses (Detmer and Chan 2007). Several consequences of impaired mitochondrial dynamics have been studied and show that mitochondria dysfunction is highly correlated with neurodegenerative diseases (Chan 2006). For instance, mutations in mitochondrial GTPase *mitofusin2* cause autosomal dominant CMT type 2A disease (Zuchner et al. 2004) and disruptions in OPA1, a protein localized at the inner mitochondrial membrane that mediates mitochondrial fusion, lead to autosomal dominant optic atrophy, an inherited form of optic nerve degeneration (Alexander et al. 2000). Moreover, recent studies have demonstrated that defects in mitochondrial function, morphology or trafficking contribute to dendritic degeneration and loss of complexity in both invertebrates and vertebrates. Mutations in the mitochondrial protein Preli-like (Prel), as well as its overexpression, cause mislocalization and fragmentation of mitochondria, leading to dendritic loss via a mechanism involving increased eIF2α phosphorylation and translational repression (Tsubouchi et al. 2009; Tsuyama et al. 2017). Moreover, the observed *prel* mutant phenotype is strikingly similar to that observed with *form3* defects. Disrupted mitochondrial distribution leads to a loss of dendritic complexity in mouse hippocampal neurons, which precedes neurodegeneration, supporting a critical role of mitochondria is stabilizing complex dendritic architectures and maintaining neuronal viability (Lopez-Domenech et al. 2016). Interestingly, defects in *prel* led to significantly reduced density of mitochondria on both CIV axons and dendrites (Tsubouchi et al. 2009); however, as *form3* mutants exhibit preferential function in stabilizing the dendritic MT cytoskeleton, we predicted that possible mitochondrial defects would be limited to the dendritic arbor, if mitochondrial trafficking was dependent upon MTs. In support of our hypothesis, we discovered that *form3* disruption inhibited dendritic trafficking of mitochondria, whereas proximal axonal traffic was unaffected, indicating an important functional role of the MT cytoskeleton in mediating mitochondrial trafficking and dynamics on the dendritic arbor. These findings are also intriguing in terms of conserved functions between Form3 and INF2, which has been demonstrated to affect mitochondrial length and ER-mitochondrial interactions (Korobova et al. 2013).

In addition to mitochondrial trafficking defects, we also postulated that defects in MT cytoarchitecture may impair trafficking of satellite Golgi on the dendritic arbor. Satellite Golgi have been shown to play important functional roles in regulating dendritic growth and branching by serving as local sites for MT nucleation to support branch extension (Ye et al. 2007; Ori-McKenney et al. 2012). Indeed, we discovered that *form3* deficient neurons also exhibited defects in satellite Golgi trafficking and in Golgi architecture indicative of fragmentation. Interestingly, previous studies have implicated INF2 in maintenance of Golgi architecture in cultured cells where disruptions in INF2 function cause Golgi fragmentation, revealing another conserved molecular function between Form3 and INF2 (Ramabhadran et al. 2011). Combined, these analyses provide important insights into the mechanistic bases of *form3-mediated* defects in dendritogenesis, and identify a critical role of a stable MT cytoskeleton in supporting the trafficking of these organelles.

### Conserved functions of form3 and INF2 in microtubule stabilization and thermosensory nociception: implications for CMT sensory neuropathy

Numerous neurological disorders including Lissencephaly, amyotrophic lateral sclerosis, spastic paraplegia, tauopathies, Alzheimer disease and CMT are linked to defects in the MT cytoskeleton and/or MT motor based transport (Franker and Hoogenraad 2013). We show *form3* is required for dendritic development, while the potential role of *INF2* is unknown with respect to neural development. Morphological defects caused by *form3* disruption can be partially rescued by expression of the INF2 FH1-FH2 domains, recovering not only lost dendritic complexity, but also MT stabilization. These analyses reveal evolutionarily conserved functions between *form3* and *INF2* in regulation of dendritic architecture and MT stabilization.

Mutations in *INF2* are known to be causative for CMT disease (Boyer et al. 2011), although the mechanistic functions of INF2 in disease pathogenesis are unclear. We observed that *form3* disruption in CIV nociceptive sensory neurons severely impairs noxious heat evoked behavioral responses resulting in peripheral insensitivity, which is consistent with sensory neuropathies observed in CMT patients. Intriguingly, we can partially revert this heat insensitivity by the introduction of INF2 FH1-FH2 in the *form3* mutant background, revealing not only morphological rescue, but also behavioral rescue. Moreover, that expression of INF2-FH1-FH2 in a *form3-IR* background can stabilize dendritic MTs and partially rescue behavioral deficits suggests that perhaps this stabilization may be more critical in restoring behavioral sensitivity than fully restoring dendritic field coverage.

Previously, CMT diseases have been characterized by progressive defects in axonal development, myelination, protein translation, and intracellular traffic of vesicles and organelles (Bucci et al. 2012; Niehues et al. 2014). CMT disease has also been linked to various defects in mitochondrial dynamics with CMT causing mutations altering energy production via a mitochondrial complex I deficiency (Cassereau et al. 2011). Our work, in addition to these mechanisms, suggests a possible additional mechanism whereby aberrant INF2 activity may lead to progressive sensory neuron dendritic atrophy which could dramatically reduce dendritic field coverage and contribute to peripheral insensitivity.

CMT sensory neuropathies manifest as impaired peripheral sensitivity to heat, cold and pain and mutations in thermosensitive TRP channels such as TRPV4 have been linked to CMT2C (Landouré et al. 2010). We discovered that *form3* mutant CIV nociceptive sensory neurons exhibit defects in dendritic trafficking of the thermosensitive TRPA channel Painless. Interestingly, previous work demonstrates that the heat sensitive TRPV1 channel is known to bind to MTs and disruption of the MT cytoskeleton leads to attenuation of TRPV1 currents impacting desensitization (Goswami et al. 2011; Lainez et al. 2010; Ferrandiz-Huertas et al. 2014). TRPV1 physically interacts with MTs and an intact MT cytoskeleton is required to preserve TRPV1 function as MT disassembly by treatment with colchicine impairs TRPV1 membrane insertion revealing a role in ion channel trafficking to the membrane (Goswami et al. 2007; Goswami 2012; Storti et al. 2012). Given that *form3* mutant neurons exhibit dendritic MT destabilization and aberrant TRP channel trafficking, these results suggest potential functional links between MT stability and TRP channel trafficking and that defects in these processes may contribute to the behavioral insensitivity to noxious heat observed in *form3* mutants.

Combined, our findings provide novel mechanistic insights into the potential etiological bases of INF2-mediated CMT sensory neuropathy, and provide evidence for functional conservation of these molecules between fly and humans. Future studies could directly investigate these potential functional links using *INF2* mutant iPSC-derived neurons to examine predicted dendritic and MT destabilization defects. The *Drosophila* model of CMT sensory neuropathy presented here provides a powerful platform for unraveling the mechanistic functions of these Formins at both the morphological and behavioral levels.

## Acknowledgements

This research was supported by NIH R01 NS086082 (DNC/GAA); NIH R15 MH086928 (DNC); R01 NS39600 (GAA); NSF BRAIN EAGER DBI-1546335 (GAA); Hungarian Brain Research Program (KTIA_NAP_13-2-2014-0007), GINOP-2.3.2-15-2016-00032 grant and the Hungarian Scientific Research Foundation (OTKA grant 109330) (JM). We acknowledge Drs. Akinao Nose, Yuh-Nung Jan Daniel P. Kiehart, Chris Q. Doe, and W. Daniel Tracey for *form3* reagents and fly strains. We thank the Bloomington *Drosophila* Stock Center (NIH P40ODO18537) and Vienna *Drosophila* Resource Center (VDRC) for fly strains used in this study.

## Movies

**Movie 1**: *form3* disruption in CIV nociceptive neurons severely impairs heat-evoked rolling behavior.

**Movie 2**: INF2-FH1-FH2 rescue of *form3* impaired heat-evoked nociceptive rolling behavior.

**Movie 3**: Optogenetic analyses implicate *form3* in sensory transduction.

## Notes

**Conflict of Interest**: The authors declare no competing financial interests

